# Age dependent impairment of home cage behavior and reactivity in *Cntnap2* knock out mouse model

**DOI:** 10.64898/2026.01.20.700626

**Authors:** Maggie Sheridan, McKenzie Rice, Ramona Mahadeshwar, Akhila Kanamarlapudi, Christina Gross, Durgesh Tiwari

**Affiliations:** Division of Neurology, Cincinnati Children’s Hospital Medical Center, Cincinnati, Ohio 45229, USA; Department of Pediatrics, University of Cincinnati College of Medicine, Cincinnati, Ohio 45229, USA; Neuroscience Graduate Program, University of Cincinnati College of Medicine, Cincinnati, OH 45267, USA

**Keywords:** epilepsy, autism, epileptogenesis, behavior, neuronal networks, neuroinflammation

## Abstract

Contactin-associated protein-like 2 (CNTNAP2) is a transmembrane protein that mediates neuron-glia interactions and regulates dendritic spine growth and neuronal migration. Mutations in the *CNTNAP2* gene are linked to autism and epilepsy. Younger *Cntnap2* KO mice mimic autism phenotypes, while older mice are a model for epilepsy. Thus, comparing behavioral phenotypes across different ages is needed to better understand the age dependent development of disordered brain networks in *Cntnap2* mutants. Male and female *Cntnap2* KO and WT controls were tested across different age groups (4, 5, 7, 9, and ∼11 months) using digging, stimulus (reactivity), and nesting assays. Older *Cntnap2* KO mice (7, 9, and ∼11 months) showed a significant increase in home cage reactivity (stimulus) assay compared to younger mice at 4 and 5 months of age. Similar trends were observed in male and female *Cntnap2* KO mice. No significant differences were observed in WT controls. A significant difference in digging assay was observed in KO female mice between younger (4 month) and older mice post nest removal. An age-dependent significant reduction in nesting behavior was observed in female KO mice; however, no difference was observed in the WT controls. Immunohistochemical analysis showed age dependent change interneuron and microglial network in *Cntnap2* KO mice. Our findings suggest disruption in home cage behavior and reactivity in older pre-epileptic *Cntnap2* KO mice indicating an age-dependent network alteration and behavior deficits.

**Significance Statement:** This study investigates the age-dependent behavioral changes in *Cntnap2* KO mice due to underlying changes in the neuronal network. It has been shown that younger *Cntnap2* KO mice display autistic behaviors and that older *Cntnap2* KO mice have epilepsy, but it is unknown how behavior is affected during the intervening period of epileptogenesis. We find that female *Cntnap2* KO mice at 11 months of age have increased reactivity and decreased motor activity compared to younger age groups, whereas WT mice show no relationship between age and behavior. Overall, the loss of *Cntnap2* alters behavior in an age-dependent and sex-specific manner, indicating progressive dysregulation of the neuronal network.

## Introduction

*CNTNAP2* plays a vital role in organizing the neuronal network and affects neuronal function. Mutations in *CNTNAP2* have been linked to neurological disorders including epilepsy [1], autism [2, 3], intellectual disability [4], and language disorders [5, 6]. CNTNAP2 is a transmembrane cell adhesion protein of the neurexin superfamily highly expressed in cortical excitatory neurons [7]. CNTNAP2 associates with calcium-calmodulin serine protein kinase, CASK, to stabilize dendritic spines during neurogenesis [8], and *Cntnap2* knockout decreases interneuron dendritic length and branching [9]. In addition, CNTNAP2 is an important mediator of both excitatory and inhibitory signaling, affecting AMPA receptor trafficking (when complexed with CASK) in excitatory neurons, [9, 10] and potassium channel distribution at the juxtaparanodes of interneurons [11].

Rodent models of *Cntnap2* reveal altered neuronal structure and behavior. Both somatosensory cortex hyperconnectivity [12] and corticothalamic hyperconnectivity [13] have been discovered in these models. *Cntnap2* KO mice have epilepsy [14, 15] and symptoms of autism, including repetitive behaviors and decreased communication [14, 16, 17] as well as social deficits [14, 18]. Evidence for the effects of *Cntnap2* on activity has been mixed, with studies showing both hypoactivity [15, 19] and hyperactivity [14, 17, 20] in mice.

Epileptogenesis refers to the formation of an excitatory network that primes the brain for epilepsy. Epileptogenesis induces behavioral changes that help clinicians identify epilepsy-prone individuals and begin therapy prior to epilepsy onset [21]. Autism is characterized by abnormal activity and sensitivity to stimuli, making it difficult to project epilepsy risk. [Altered neuronal networks predispose *Cntnap2* knockout mice to autism and epilepsy from an early age, with mice 8-14 weeks displaying autistic behaviors and increased seizure susceptibility due to reticulothalamic neuron hyperexcitability {Jang, 2025 #133}. However, the age-dependent behavioral phenotype is unknown. This study investigated age-dependent changes in home cage behavior using 4–12-month-old *Cntnap2* KO mice. We found that female KO mice at 11-12 months of age were more reactive and less active than mice at younger ages, whereas no significant age-dependent differences were observed in wild type mice. We also found that KO mice had larger interneurons size and microglia, which mark network remodeling and neuroinflammation. Overall, these findings show that loss of *Cntnap2* and subsequent network dysregulation leads to age-dependent and sex-specific alterations in various home cage reactivity behaviors, which may represent a key stage for intervention in epilepsy and shed light on common mechanisms of this behavior during epileptogenesis.

## Material and Methods

### Animals

All animal procedures were approved by the Institutional Animal Care and Use Committee of CCHMC and complied with the Guideline for the Care and Use of Laboratory Animals. *Cntnap2^tlacz/tlacz^* mice WT mice were initially procured from The Jackson Laboratory (Bar Harbor, ME, USA), cat. no. 017482, RRID: IMSR_JAX:017482 and 000664, RRID: IMSR_JAX:000664 respectively. These mice were set up for breeding generated using *Cntnap2* KO x *Cntnap2* KO and WT x WT breeding pairs in our mouse vivarium facility at CCHMC. Genotyping was performed for the *Cntnap2* mice as per the previously established protocol from Jackson laboratory. Pups were weaned at P21–28 and were housed with same sex littermates (maximum 4 per cage) in a standard cage with food and water provided *ad libitum.* Mice were used after weaning at various time points as indicated in respective experimental sections. Mice were maintained on a 14:10 hour light: dark cycle, and all experiments were performed during the light cycle.

### Behavioral experiments

For all experiments, mice were individually housed. The experiments were conducted in the order of least to most chronic behavior: first nesting, then digging, then stimulus assay as per the timeline (see **Figure 1A**).

**Figure 1.**
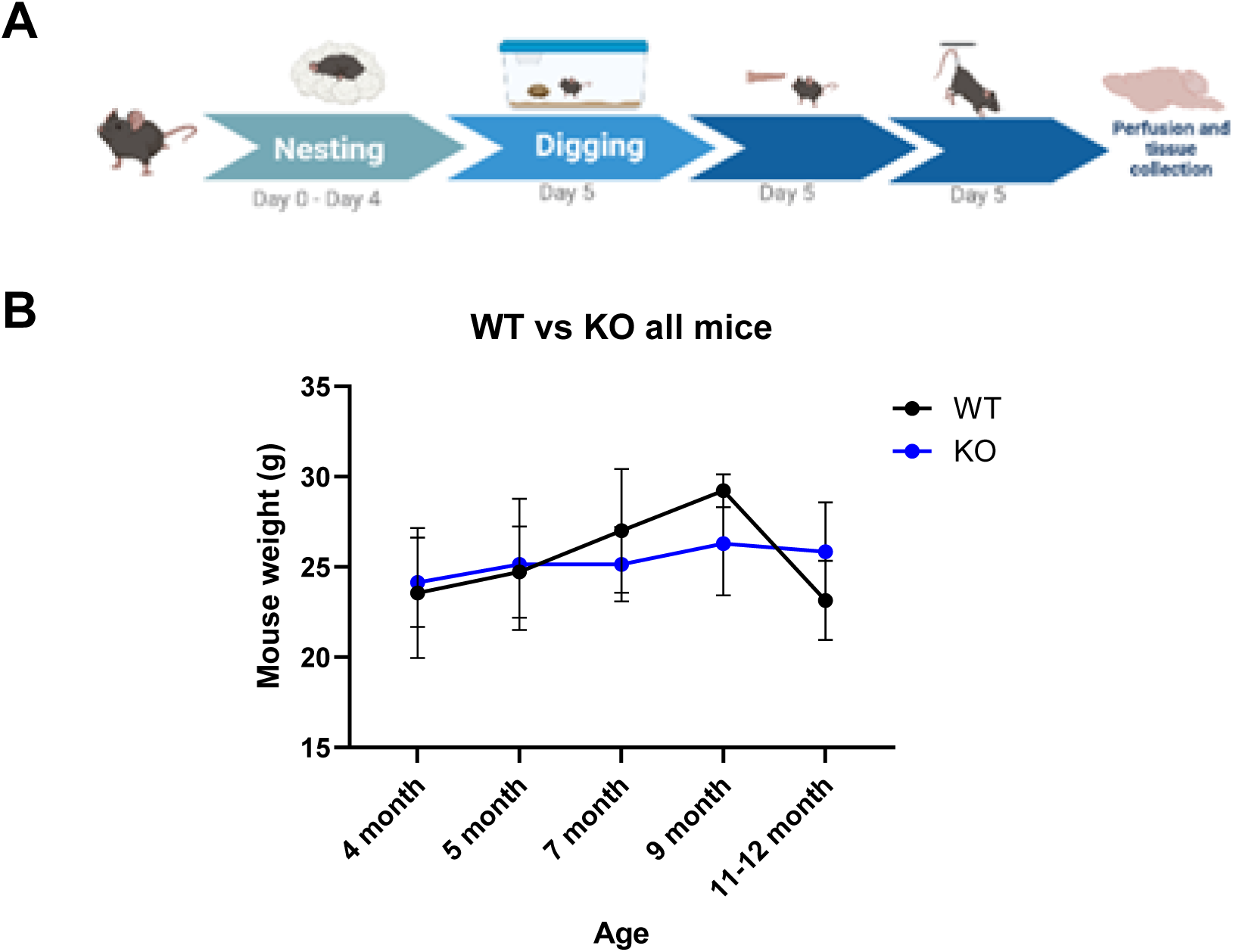
(A) Timeline of experiment. (B) Mouse weights. WT and KO mice did not have significantly different weights at any age.

#### Nesting

Mice were initially separated and single housed for 3 days before the assay. After a 3-day acclimation period, nesting behavior was assessed at 2 hr and 24 hr post as per the described protocols [22, 23]. Briefly, nests were scored on a rating scale of 1 to 5. A score of 1 indicates more than 90% intact nestlet, 2 indicates 50–90% intact nestlet, 3 indicates >90% nestlet torn with no identifiable nest, 4 indicates >90% nestlet torn with an identifiable nest, and 5 indicates >90% nestlet torn with a nest whose walls are taller than the mouse. The leftover nestlet was weighed and the percentage torn was calculated and compared. While scoring based on visual observation is less objective than weighing used nestlet, it allows for evaluation of the nest regardless of how much nestlet is used.

#### Digging Behavior

The digging assay was performed one day after the nesting. Water bottle, cage lid and metal food top were removed to avoid disruption and ensure visibility and mice were initially habituated for 10-minute duration. Post habituation, time spent digging was recorded for each mouse during a 10-minute period with the nestlet in the cage. After 10 minutes, the nestlet was removed from the cage and the time spent digging was recorded for another 10-minute period. The digging behavior was analyzed as total time spent digging with no nestlets.

#### Stimulus

Stimulus assays were performed 10 minutes after the digging assay. An experimenter applied 1) a light puff of air to the back, 2) light pressure to the base of the tail, 3) light pressure to the back, and 4) light pressure to the head. The stimuli were applied in the same order every time. Responses were scored in 0.5 increments from 0 (no reaction) to 3 (most reaction) using criteria from Calsbeek et al, 2021. All response scores were summed to obtain a total reactivity score. After the four stimuli, mice were suspended by the tail for two minutes, returned to the cage, and monitored for seizures.

#### Tail suspension

Tail suspension assay was used to evaluate depressive behavior in the mice. Mice were suspended by the tail by an experimenter six inches above the cage floor for two minutes. A smaller time period was chosen to avoid the occurrence of any seizure behavior especially in older mice as reported in previous study (Ref) and we observed no seizure during handling our paradigm allowing us to study the paradigm. The video recordings were monitored and analyzed for onset and number of freezing instances and latency to freezing were recorded.

### Immunohistochemistry

Mice were injected with pentobarbital (equivalent to body wight) and later perfused with PBS using trans cardiac perfusion followed by 4% PFA. Brains were extracted and then cryoprotected with sucrose (10% and 30%, 24 hours each) followed by freezing using dry ice and liquid nitrogen and stored in -80 until sectioning. 25µm PFA-fixed brain sections were processed using cryostat and were first stained for primary antibodies (GFAP (glial) (cat. # ab7260, RRID:AB_ AB_305808, *Abcam*, Cambridge, UK, 1:500), Iba1 ((cat. # ab7260, RRID:AB_2636859, *Abcam*, Cambridge, UK, 1:500) and parvalbumin (inhibitory neurons)) (cat. # ab32895, RRID: AB_777105, *Abcam*, Cambridge, UK, 1:500) along with equivalent secondary antibodies Alexa Fluor 488 donkey anti-rabbit (1:500; Jackson ImmunoResearch; RRID:AB_2340619) and Alexa Fluor 647 donkey anti-mouse ((1:500; Jackson ImmunoResearch; RRID:AB_2340862)), followed by DAPI nuclei counterstain (1:5000, sigma Aldrich D9542-1mg; RRRID:AB_2869624). Sections were imaged with a BAF Widefield Nikon NiE Upright #1 microscope at 4x, 10x, and 40x magnification. Images were captured with the Nikon Elements 360 software. The images were imported to trace was drawn over the sub hippocampal regions using mouse brain atlas as a reference and counted manually. A cell count was performed at least 3 mice per group with sections collected from anterior and posterior regions with in stated bregma level -1.34 to -2.54.

### Statistical Analyses

Statistical Analyses was performed using GraphPad Prism v10.5.0 (*GraphPad,* San Diego, CA, USA), SPSS (*IBM*, Armonk, NY, USA) or R software. The data was analyzed for normality using the Shapiro-Wilk test and parametric or nonparametric. Statistical tests were performed as appropriate and are mentioned in the individual result section or figure legends along with sample size and the data were screened for outliers using the ROUT method (Robust Regression and Outlier Removal*, GraphPad*). Error bars represent mean +/-standard error of the mean, and dots are individual mice.

## Results

### Body weight does not differ between WT and *Cntnap2* KO mice

All mice studied were initially screened for their body weight to compare any effect of genotype on overall development which could potentially interest the outcome. The body weight was measured at the start of the study and then were individually housed. No significant difference was observed between WT and KO mice in any age group (4, 5, 7, 9, and 11-12 months) (**Figure 1B**) indicating no effect of genotype on development.

### Aging decreases nesting behavior in *Cntnap2* KO female mice

Nesting behavior is a common home cage behavior used to study cognitive function and evolutionary conserved development and in mice (Deacon et al 2006). Nesting behavior was first performed at 2hr and 24 period to examine how aging affects nesting at different timepoints while mice undergo epileptogenesis. After 2 hours, no significant differences were observed in nesting score between age groups of either KO or WT mice (**Figure 2B, C**). Further, no significance was found when male mice only (**Figure 2B.1, C.1**), and female mice only (**Figure 2B.2, C.2**) were analyzed. Moreover, at 24 hours timepoints no significant age-dependent differences were observed in nesting score (**Figure 2D**) or percentage nestlet torn ***(see supplementary figure S1)***, when all KO mice were analyzed. However, when female KO mice only were analyzed, mice 11-12 months of age had a significantly lower nesting score than mice 8-9 months (p = 0.033), 5 months (p = 0.016), and 4 months (p = 0.025) of age **(Figure D.2**). These findings suggests that *Cntnap2* knockout affects the neuronal network of females differently than males. In addition, no age-dependent significant differences were observed in WT mice (**Figure 2E**). As mice age, the loss of *Cntnap2* leads to abnormal network development and a decrease in motor activity in females.

**Figure 2.**
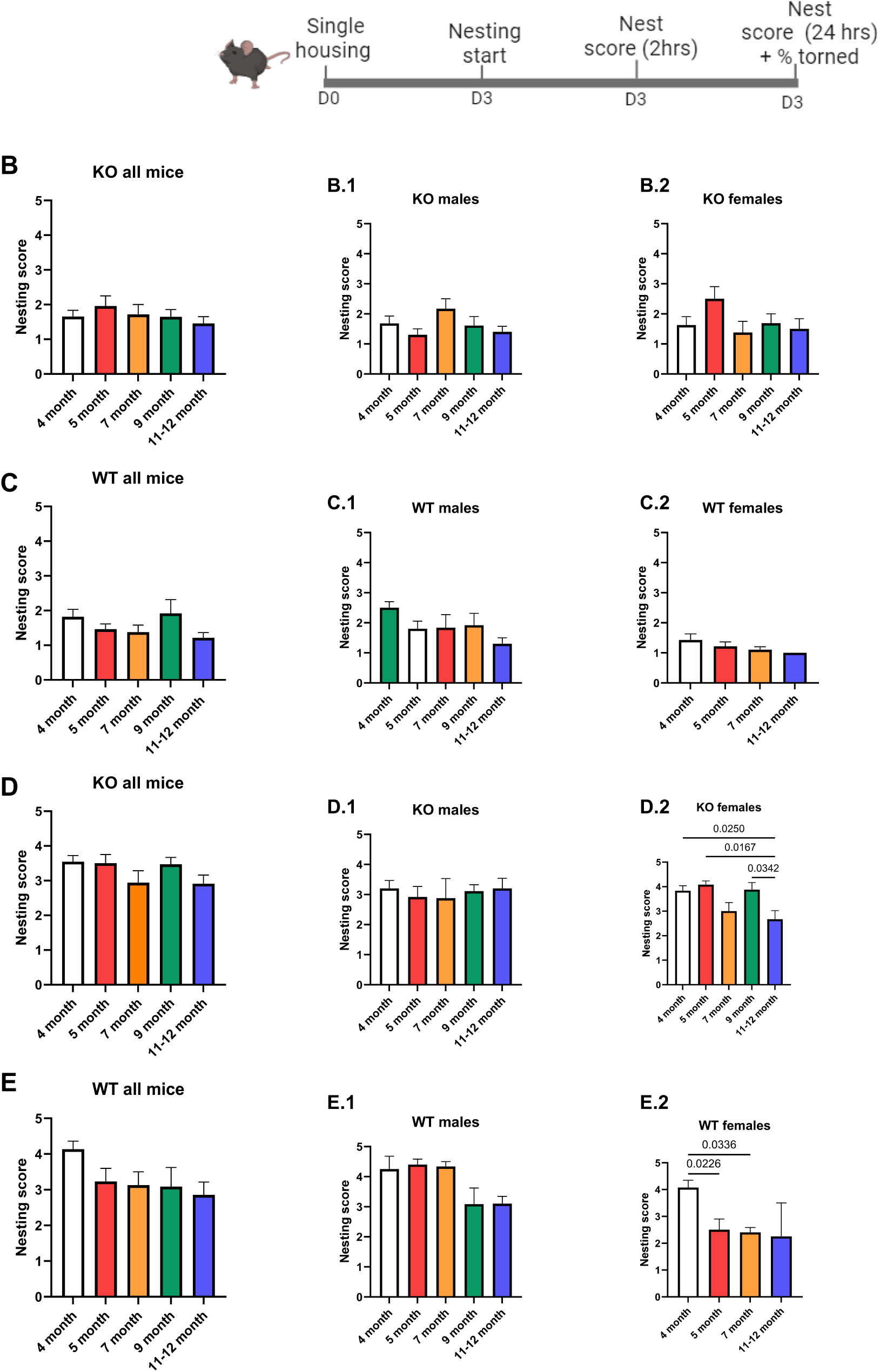
(A) Timeline for nesting and score recording. 2-hour nesting: (B, C) Neither KO nor WT mice showed a significant relationship between nesting score and age after 2 hours (p>0.05) (one-way ANOVA, n=3-23). 24-hour nesting: (D) All KO mice and male mice (D.1) did not have significantly different nesting scores after 24 hours (p>0.05), but female mice (D.2) did (**p=0.0065) (one-way ANOVA, n=4-12). 11–12-month-old female mice had significantly lower nesting scores than the 9 month (*p=0.0342), 5 month (*p=0.0167), and 4-month groups (*p=0.0250) (Tukey’s test, n = 6-12). (E) Wild type at different ages did not have significantly different nesting scores after 24 hours, and neither male (E.1) nor female (E.2) mice alone had significantly different nesting scores between age groups (one-way ANOVA, n=3-12).

### Aging decreases digging in *Cntnap2* KO female mice

Digging assay is commonly assessed to test anxiety and repetitive behavior which is a feature of neurological disorders such as epilepsy and autism {Pond, 2021 #355}. When male and female *Cntnap2* KO mice were combined, an age-dependent decrease in digging was observed (p = 0.0197) (**Figure 3B**). However, this relationship was driven more by female than by male mice. When female mice alone were analyzed, the decrease in time spent digging at older ages showed a trend towards approaching significance ((p = 0.062) (**Figure 3B.2**); male mice alone did not show any significance between age groups (**Figure 3B.1**). No age-dependent differences were observed in wild type mice (**Figure 3C**) and no effect of sex was also observed (**Figure 3C.1 and 3C.1).** These findings reinforces that loss of *Cntnap2* exerts a differential effect on the neuronal networks in male and female mice, thus indicating that loss of *Cntnap2* causes an abnormal decrease in motor activity and anxiety behavior.

**Figure 3.**
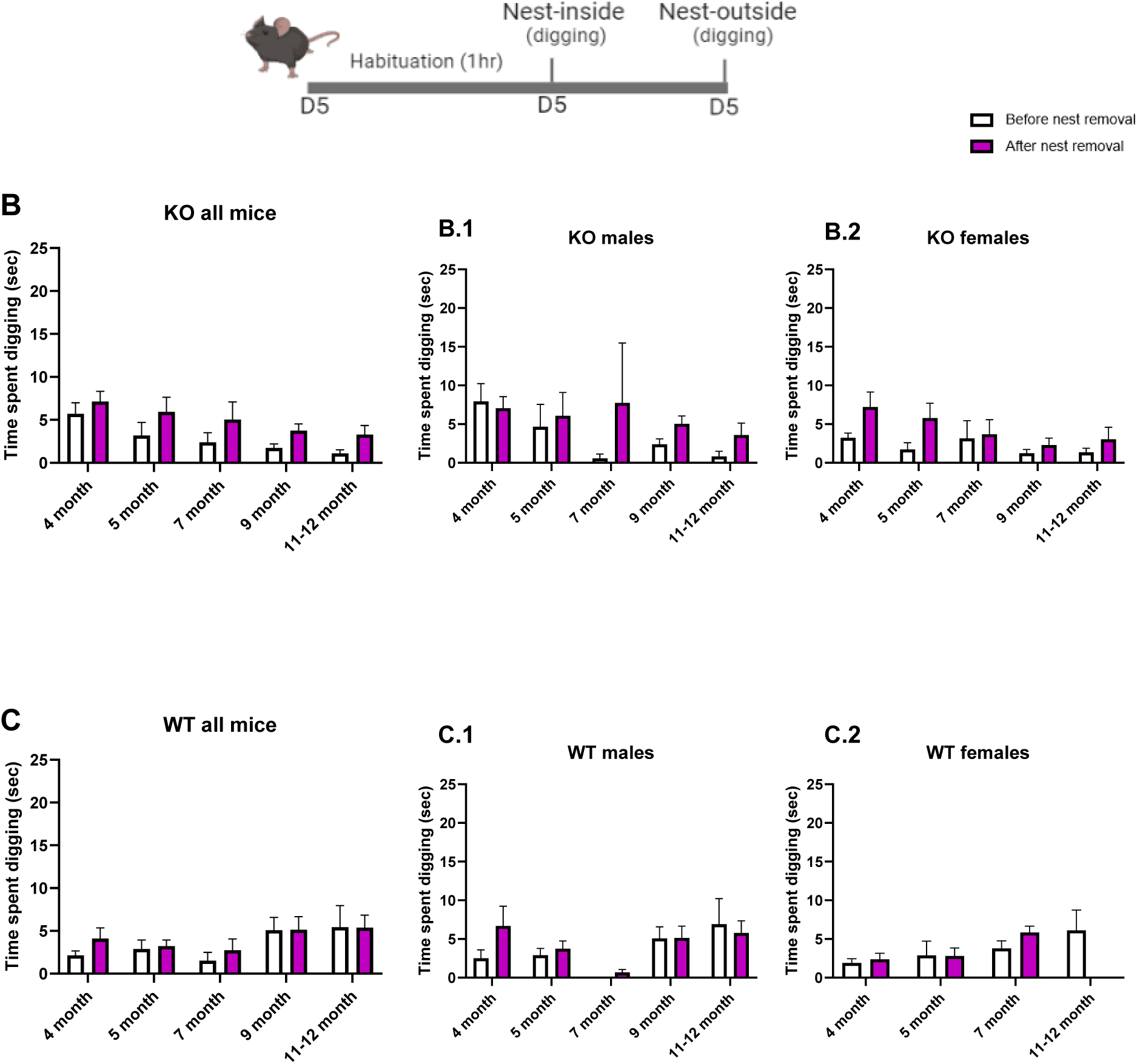
(A) Timeline for digging behavior. (B) Cntnap2-KO mice displayed a significant age-dependent decrease in time spent digging (two-way ANOVA, *p=0.0197, n=7-23). Male mice (B.1) had a nonsignificant age-dependent decrease (two-way ANOVA, p=0.2755, n=2-12), and female mice (B.2) had a nearly significant age-dependent decrease (two-way ANOVA, p=0.0625, n=3-11). (C) There was no significant effect of age on time spent digging for all WT mice, male mice (C.1), or female mice (C.2) (all p>0.05) (one-way ANOVA, n=2-23).

### Aging increases stimulus reactivity in both male and female *Cntnap2* KO mice

An elevated sensory response or hyper-reactivity to an external stimulus is observed due to an imbalance of excitation and inhibition in ASD and during epileptogenesis (Green et al 2013). Hence, we studied the response to stimuli within the home cage in the cntnap2 KO mice. Our findings indicated that *Cntnap2* KO mice at 11-12 months of age displayed a significantly increased reactivity score compared to all younger age groups: 4 months (p = 0.003), 5 months (p = 0.025), 7 months (p = 0.0003), and 8-9 months (p = 0.005) (**Figure 4B**). The age dependence of reactivity differed by sex. For females, 11–12-month-old mice had a significantly greater reactivity score than 7-month-old (p = 0.017) and 4-month-old mice (p = 0.015) (**Figure 4B.2**). For males, 11–12-month-old mice had a significantly greater reactivity score than 7-month-old mice (p = 0.023) (**Figure 4B.1**). While female WT mice did not show any significant differences in reactivity between age groups (**Figure 4C.2**), younger male WT mice showed greater reactivity: 5-month-old male WT mice had a significantly higher reactivity score than 8–9-month-old mice (**Figure 4C.1**). *Cntnap2* KO can be clearly linked to increased reactivity in older female mice, but the effect on males is more ambiguous, given that KO males in the oldest age group displayed increased reactivity whereas WT males in the youngest age group displayed increased reactivity. In all groups except WT males, 7-month-old mice had the lowest reactivity score and 10–12-month-old mice had the highest reactivity score, suggesting an initial decrease in reactivity is followed by an increase. Dysregulation of the neuronal network after *Cntnap2* knockout elevates reactivity in older male and female mice in age dependent manner indicating a progressive network disruption in the model.

**Figure 4.**
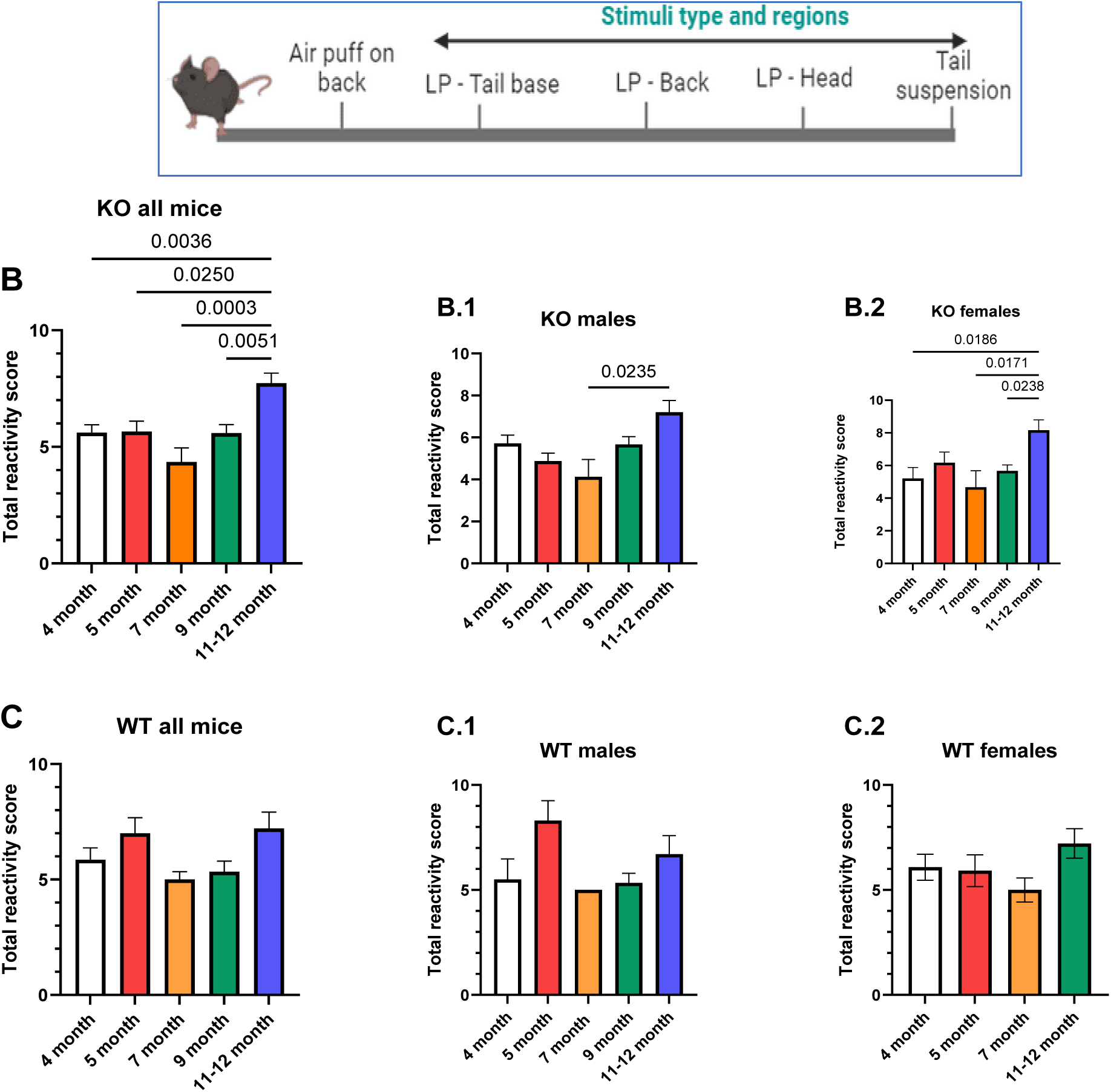
(A) Timeline for stimulus behavior. (B) A significant effect of age on reactivity behavior (average score) was observed in Cntnap2 KO mice, where mice 11-12 months of age were significantly more reactive than 9 month (**p=0.0051), 7 month (**p=0.0036), 5 month (*p=0.0250), and 4-month-old mice (**p=0.0036) (Tukey’s test, n=7-23). For male mice analysis, mice 11-12 months old were significantly more reactive than mice 7 months old (*p=0.0235) (Tukey’s test, n=5,4). For female mice 11-12 months old were significantly more reactive than mice 8-9 (*p=0.0238), 7 (*p=0.0171), and 4 months old (*p=0.0186) (Tukey’s test, n=3-9). (B) Overall, no significant difference in home cage reactivity was observed for WT controls however, a trend toward significance (p=0.0931) was driven by male mice (*p=0.0117) but not female mice (p=0.1651) (one-way ANOVA, n=2-11). 5-month-old male mice were significantly more reactive than the 7-month (*p=0.0290) and 8–9-month (*p=0.0187) groups (Tukey’s test, n=5,3,6) indicating an effect of sex on home cage reactivity in the WT mice.

### Aging increases handling-induced freezing in *Cntnap2* KO mice

Tail suspension assay was used to analyze the effect of age on depressive like behavior in *Cntnap2* KO model. When all *Cntnap2* mice were analyzed without differentiating by sex, 9-month-old mice had significantly more freezing episodes (p = 0.0177) (**Figure 5C**) and a trend toward earlier freezing onset (p = 0.1090) than 7-month-old mice (**Figure 5E**). For male mice, 9-month-old mice had significantly more freezing episodes than 7-month-old mice **(see supplementary section, figure S2C**) and 5- and 9-month-old mice had significantly earlier freezing onset than 7-month-old mice **(see supplementary figure S2G**); however, female mice did not show any significant differences in either freezing episodes or freezing onset **(see supplementary figure S2D, H)**. In addition, WT mice showed no age-dependent differences in freezing episodes or freezing onset **(Figure S2A, B, E, F)**. No handling-induced seizures were observed. These findings indicate an age specific effect of increased freezing/depressive behavior in *Cntnap2* mice due to potentially altered neuronal network due to epileptogenesis.

**Figure 5.**
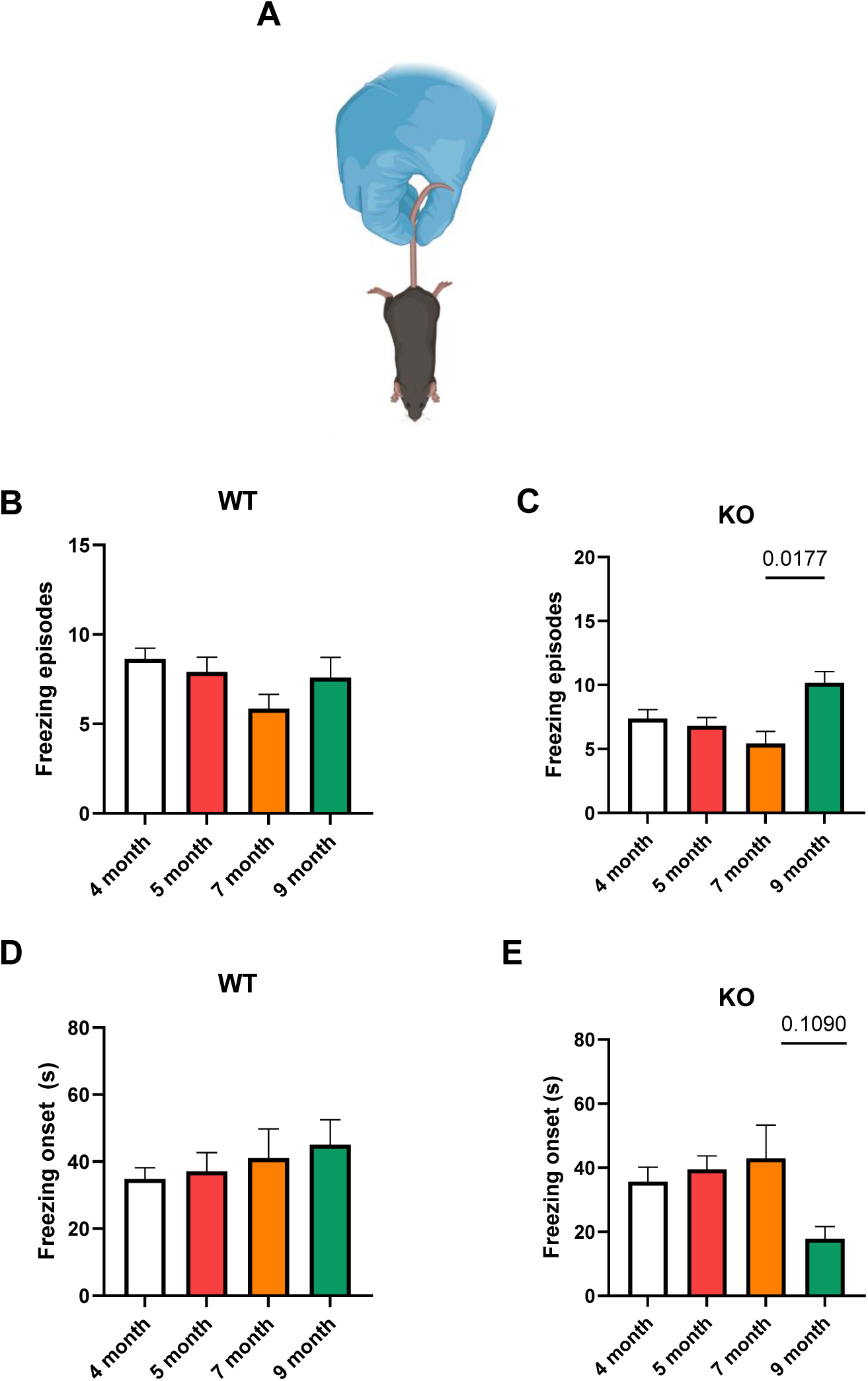
(A) Schematic of tail suspension. (B) WT mice did not display significant differences in number of freezing episodes between any age groups (Tukey’s test, p > 0.05, n=5-11). (C) 9-month-old KO mice had a significantly greater number of freezing episodes than 7-month-old mice (Tukey’s test, *p=0.0177, n=7, 6). (D) WT mice did not display significant differences in freezing onset between any age groups (Tukey’s test, p > 0.05, n=5-11). (E) 9-month-old KO mice had a trend toward an earlier freezing onset than 7-month-old mice (Tukey’s test, p = 0.109, n=7, 6).

### *Cntnap2* KO mice show altered cell morphology

Reduced interneuron function and increase in excitation and neuroinflammation is commonly observed in Cntnap2 KO mouse however hasn’t been analyzed in age specific manner. Interneuron and inflammatory-microglial network were studied by means of immunostaining using Albumin and Iba1 antibodies in mouse brain hippocampal region (CA1, dentate gyrus and Hilus regions) at 3 timepoints (4 months, 9months and 11 months) to study network development over time in *Cntnap2* KO mice compared to WT controls. Quantitative analysis showed no difference in the number of hippocampal interneuron population (parvalbumin staining) between WT and KO mice in CA1 or DG (**Figure 6A and 6B**). Similarly, there is no difference in microglial population (Iba1 staining) between WT and KO mice in CA1 or DG (**Figure 7A and 7B**)., potentially due to a smaller sample size. However, qualitative morphological analysis has shown an increased cell body in parvalbumin with larger interneuron and bigger-proliferated microglia with more branching in *Cntnap2* KO mice as compared to WT mice **(see supplementary figure S3)**. B These results indicated an altered morphology of interneuron and increased neuroinflammation in the cntnap2 model indicating an altered neuronal network potentially correlating with our behavioral findings.

**Figure 6:**
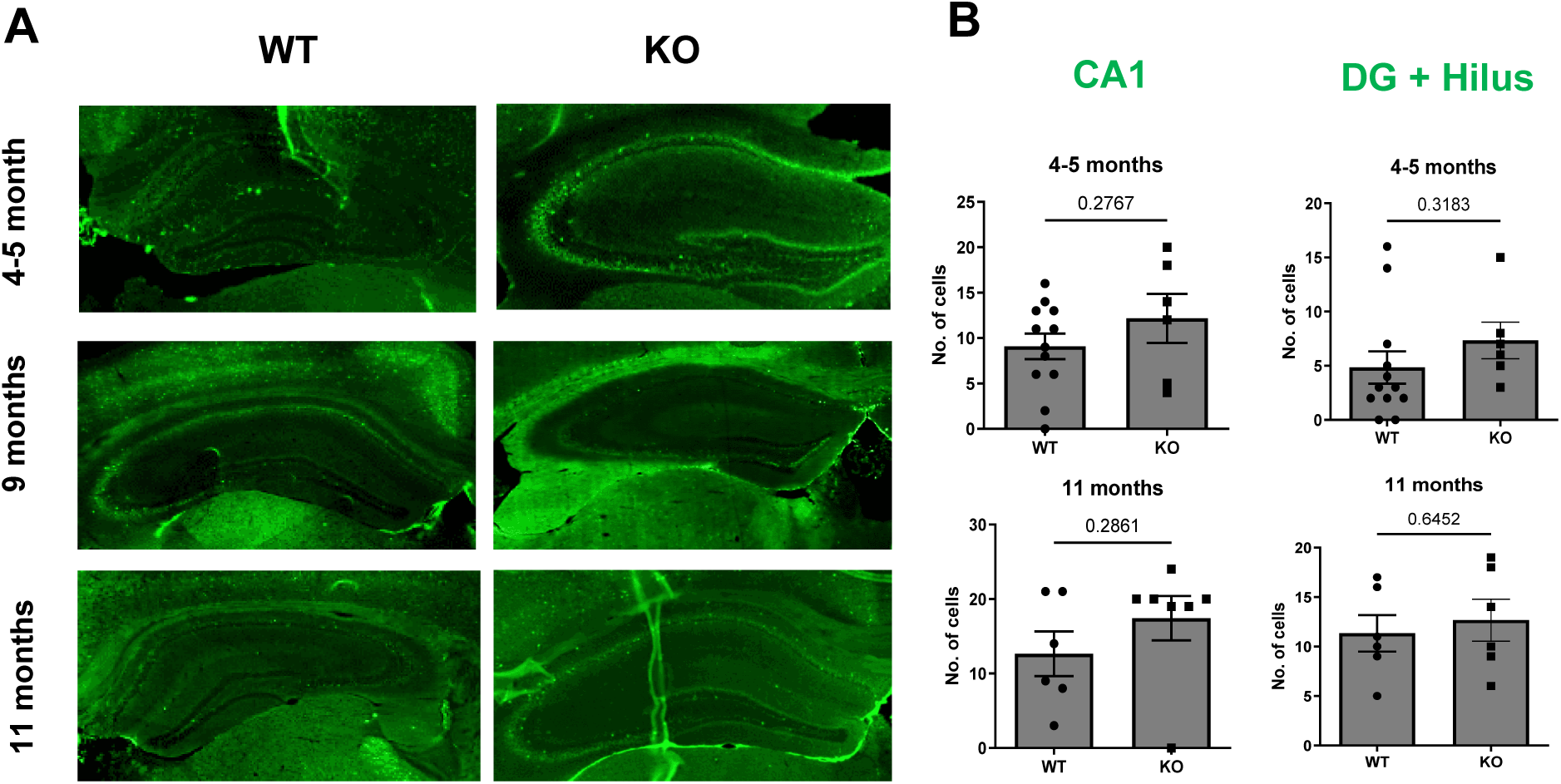
(A) Representative image of parvalbumin (interneuron) immunostaining at 4–5, 9 and 11 -month-old *Cntnap2* mice and WT controls. Quantitative analysis showed no significant difference in the number of parvalbumin cells (p values shown on the graphs) in the CA1 (B) and DG+Hilus region in *Cntnap2 KO* mice compared to the WT controls. Error bars represent SEM.

**Figure 7:**
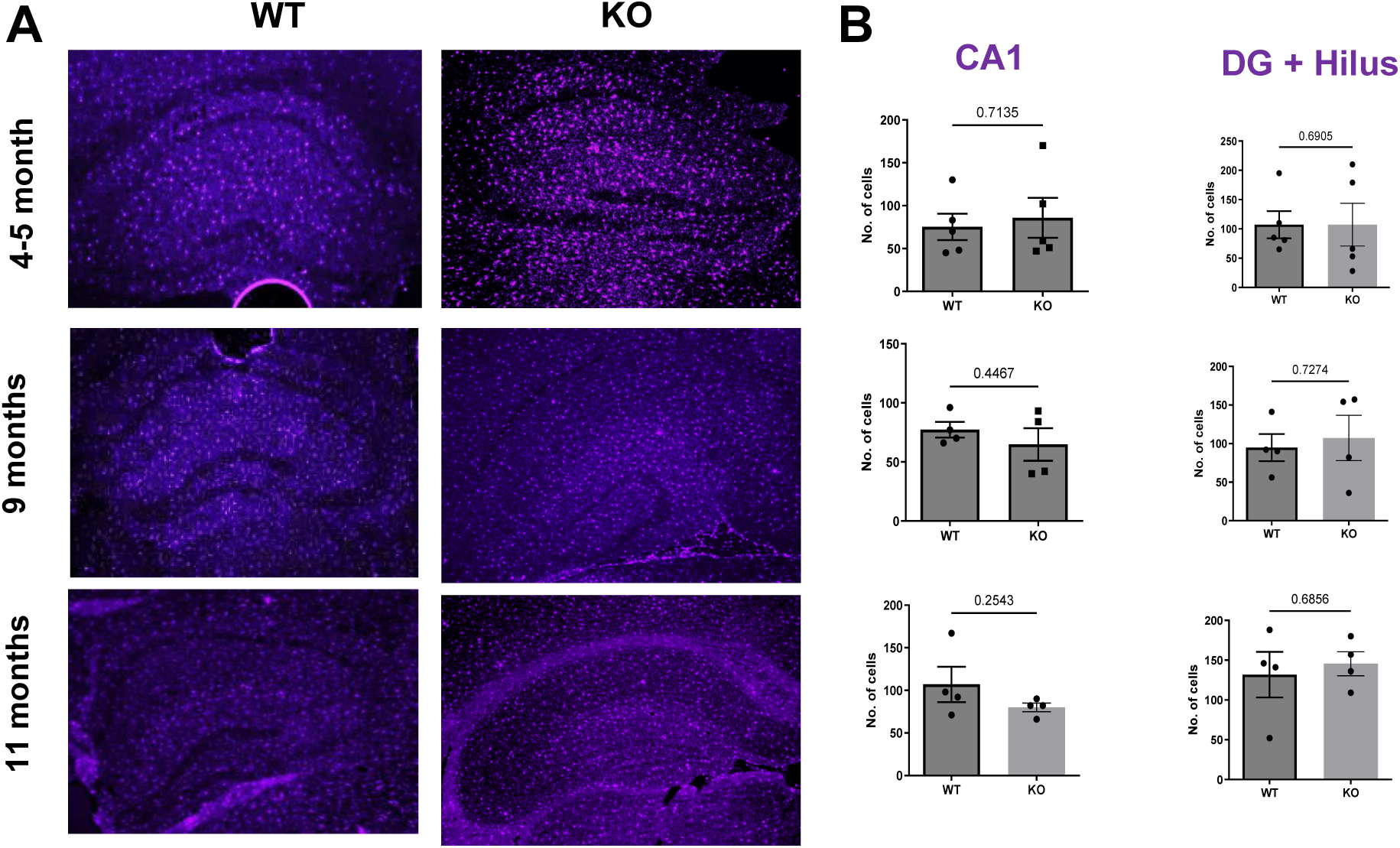
(A) Representative image of Iba1 (microglial) immunostaining at 4–5, 9 and 11 -month-old *Cntnap2* mice and WT controls. Quantitative analysis showed no significant difference in the number of Iba1 positive cells (p values shown on the graphs) in the CA1 (B) and DG+Hilus region in *Cntnap2 KO* mice compared to the WT controls. Error bars represent SEM.

## Discussion

This study shows that deletion of Cntnap2 leads to age-dependent home cage behavior and reactivity in *Cntnap2* KO mice potentially during the period of epileptogenesis progressing from autism to an epileptic brain circuit. Our battery of behavior test showed deficits at various timepoints studied however most deficits was observed at 11–12-month-old *Cntnap2* KO mice as compared to WT control group. In addition, behavioral trends differ based on the type of behavior. Nesting and reactivity generally reached a low at seven months of age before increasing through 11-12 months of age, digging decreased across all age groups, and freezing increased at 9 months of age (males only). Since these behaviors are encoded in different neuronal circuits, which are regulated differently the finding indicated disruption in one circuit over the period of time in age specific manner. Overall, this pattern of progressive behavioral deficits during the period of epilepsy development (epileptogenesis) aligns with clinical and preclinical studies indicating an alteration in home cage behavior during development (Kwon et al., 2019).

Autism is thought to have a male-enhanced phenotype [19]. While our study did not investigate traditional autistic phenotypes between males and females, we found a greater change in home cage behavior in females in three of four assays. This indicates that neuronal network organization differs by sex, which may contribute to differential disease phenotypes. Moreover, this work also adds to the growing literature evidence on significant effect of sex in network regulation during epileptogenesis {Reddy, 2021 #356{Pais, 2025 #358}}.

Neither WT nor *Cntnap2* KO mice were epileptic or showed any behavioral seizures in any age group however the time tested represents a period of progressive epileptogenesis. Our previous unpublished studies have also shown potential electrographic seizure around 14 months of age indicating the period tested to be of epileptogenesis. In out behavioral findings a progressive disruption of behavior was also observed as the *Cntnap2* KO mice approached 10-11 months of age, significant changes in nesting, digging, and reactivity relative to younger ages were measured, but no changes were measured prior to this time. Therefore, 10 months represents a critical period where neuronal network dysregulation begins to affect behavior. In particular, the increase in reactivity supports a hyperexcitable network on the cusp of epilepsy.

There is limited literature exploring the home cage reactivity in autism and epilepsy mouse model [19, 32]. One of the previous studies by *Angelakos et al* 2019 have utilizing home cage activity in various *Shank3b* KO*, Cntnap2* KO*, Pcdh10-het, and Fmr1* KO mouse model of autism highlighted alteration over a shot 24-day period [19]. However, our study comprehensively analyzes one of the main *Cntnap2* KO mouse model in age specific manner over various ages points indicating behavioral impairment and its progression over time. Angelakos et al 2019 also showed a sex specific effect in the *Cntnap2* KO model which was also observed in our hands. However, the previous study utilized an entirely different set of behavioral paradigms whereas our choice behavior was based on home cage behavior and reactivity in a familiar environment.

We observed no changes in cell count for interneurons (parvalbumin staining) and neuroinflammation-microglia (Iba1 staining) in the hippocampal brain regions. However, *Cntnap2* KO mice had larger interneurons, which could indicate a compensatory process to hold the hyperexcitable network in check. Further, in epileptic mice, hyperactive interneurons activate microglia through GABAergic signaling [24]. Though *Cntnap2* KO mice were not epileptic, they showed evidence of microglial activation, as they had larger and less branched microglia than WT mice. These findings indicated interneuron activation and neuroinflammation, which is critical in the development of an epileptic network [25].

One limitation of this study is that hyperreactivity was only measured with respect to mechanical stimuli. It is possible that the mice could react differently to thermal, visual, auditory, or taste stimuli. Another limitation is that motor activity was assessed via digging, which may differ from other types of activity such as walking, rearing, and exploring. Further, observing motor activity over multiple days would help to establish the chronic nature of the phenotype. The n numbers can also be attributed to limitation in result interpretation at certain timepoints hence our findings were limited to the number of animal studies. Additionally, another limitation which could indicate mice were only studied up to 12 months of age.

In summary, this study showed age-dependent changes in home-cage behavior in *Cntnap2* KO mice. When combined with the hyperexcitable network, these behaviors are an additional marker of epileptogenesis. The mice’s transition from an autistic to an epileptic network indicates that autism and epilepsy pathogenesis are related. Unlike in other studies, *Cntnap2* KO mice did not exhibit motor stereotypies [14, 26] or excessive hyperactivity [14, 18, 26]; rather, they had decreased instinctive behaviors such as digging, stimulus reactivity and nesting. Meanwhile, the hyperreactivity phenotype is common to both other rodent models [26] and young human patients with autism [27, 28]. Future studies will involve other tests to assess anxiety and social behavior and include older age groups. Neuroimaging can be used to see how not just the components but the structure of the network changes over time. Characterizing the onset, directionality, and magnitude of behavioral changes during epileptogenesis may thus help to identify specific dysregulated neuronal circuits and associated behavior which can potentially assist in early detection of epilepsy.

## Author Contributions

MS Analyzed data, Wrote the paper

MR Performed research, Training

RM Performed research, Analyzed data

AK Performed research, Analyzed data

CG Result discussions and data interpretation

DT Designed and supervised research, performed research, analyzed data, Wrote the paper

## Acknowledgements

Thanks to all Tiwari and Gross lab members.

## References

1. Leonardi, E., et al., CNTNAP2 mutations and autosomal dominant epilepsy with auditory features. Epilepsy Res, 2018. 139: p. 51–53.

2. Alarcón, M., et al., Linkage, association, and gene-expression analyses identify CNTNAP2 as an autism-susceptibility gene. Am J Hum Genet, 2008. 82(1): p. 150–9.

3. Arking, D.E., et al., A common genetic variant in the neurexin superfamily member CNTNAP2 increases familial risk of autism. Am J Hum Genet, 2008. 82(1): p. 160–4.

4. Shiota, Y., et al., A common variant of CNTNAP2 is associated with sub-threshold autistic traits and intellectual disability. PLoS One, 2021. 16(12): p. e0260548.

5. Vernes, S.C., et al., A functional genetic link between distinct developmental language disorders. N Engl J Med, 2008. 359(22): p. 2337–45.

6. Peter, B., et al., Replication of CNTNAP2 association with nonword repetition and support for FOXP2 association with timed reading and motor activities in a dyslexia family sample. J Neurodev Disord, 2011. 3(1): p. 39–49.

7. St George-Hyslop, F., et al., Loss of CNTNAP2 Alters Human Cortical Excitatory Neuron Differentiation and Neural Network Development. Biol Psychiatry, 2023. 94(10): p. 780–791.

8. Gdalyahu, A., et al., The Autism Related Protein Contactin-Associated Protein-Like 2 (CNTNAP2) Stabilizes New Spines: An In Vivo Mouse Study. PLoS One, 2015. 10(5): p. e0125633.

9. Gao, R., et al., The CNTNAP2-CASK complex modulates GluA1 subcellular distribution in interneurons. Neurosci Lett, 2019. 701: p. 92–99.

10. Varea, O., et al., Synaptic abnormalities and cytoplasmic glutamate receptor aggregates in contactin associated protein-like 2/Caspr2 knockout neurons. Proc Natl Acad Sci U S A, 2015. 112(19): p. 6176–81.

11. Poliak, S., et al., Caspr2, a new member of the neurexin superfamily, is localized at the juxtaparanodes of myelinated axons and associates with K+ channels. Neuron, 1999. 24(4): p. 1037–47.

12. Balasco, L., et al., Somatosensory cortex hyperconnectivity and impaired whisker-dependent responses in Cntnap2(-/-) mice. Neurobiol Dis, 2022. 169: p. 105742.

13. Gandhi, T., et al., Neuroanatomical Alterations in the CNTNAP2 Mouse Model of Autism Spectrum Disorder. Brain Sci, 2023. 13(6).

14. Peñagarikano, O., et al., Absence of CNTNAP2 leads to epilepsy, neuronal migration abnormalities, and core autism-related deficits. Cell, 2011. 147(1): p. 235–46.

15. Thomas, A.M., et al., Cntnap2 Knockout Rats and Mice Exhibit Epileptiform Activity and Abnormal Sleep-Wake Physiology. Sleep, 2017. 40(1).

16. Zhang, Q., et al., Contactin-associated protein-like 2 (CNTNAP2) mutations impair the essential α-secretase cleavages, leading to autism-like phenotypes. Signal Transduct Target Ther, 2024. 9(1): p. 51.

17. Jang, S.S., F. Takahashi, and J.R. Huguenard, Reticular thalamic hyperexcitability drives autism spectrum disorder behaviors in the Cntnap2 model of autism. Sci Adv, 2025. 11(34): p. eadw4682.

18. Selimbeyoglu, A., et al., Modulation of prefrontal cortex excitation/inhibition balance rescues social behavior in CNTNAP2-deficient mice. Sci Transl Med, 2017. 9(401).

19. Angelakos, C.C., et al., Home-cage hypoactivity in mouse genetic models of autism spectrum disorder. Neurobiol Learn Mem, 2019. 165: p. 107000.

20. Gandhi, T. and C.C. Lee, Behavioral Physiology of the CNTNAP2 Knockout Mouse. HSOA Trends Anat Physiol, 2025. 6.

21. Terrone, G., et al., Preventing epileptogenesis: A realistic goal? Pharmacol Res, 2016. 110: p. 96–100.

22. Deacon, R.M., Assessing nest building in mice. Nat Protoc, 2006. 1(3): p. 1117–9.

23. Gross, C., et al., Increased expression of the PI3K enhancer PIKE mediates deficits in synaptic plasticity and behavior in fragile X syndrome. Cell Rep, 2015. 11(5): p. 727–36.

24. Chen, Z.P., et al., GABA-dependent microglial elimination of inhibitory synapses underlies neuronal hyperexcitability in epilepsy. Nat Neurosci, 2025. 28(7): p. 1404–1417.

25. Sanz, P., T. Rubio, and M.A. Garcia-Gimeno, Neuroinflammation and Epilepsy: From Pathophysiology to Therapies Based on Repurposing Drugs. Int J Mol Sci, 2024. 25(8).

26. Scott, K.E., et al., Loss of Cntnap2 in the Rat Causes Autism-Related Alterations in Social Interactions, Stereotypic Behavior, and Sensory Processing. Autism Res, 2020. 13(10): p. 1698–1717.

27. Posar, A. and P. Visconti, Sensory abnormalities in children with autism spectrum disorder. J Pediatr (Rio J), 2018. 94(4): p. 342–350.

28. Niedźwiecka, A. and E. Pisula, Symptoms of Autism Spectrum Disorders Measured by the Qualitative Checklist for Autism in Toddlers in a Large Sample of Polish Toddlers. Int J Environ Res Public Health, 2022. 19(5).

29 Kwon OY, Park SP. Depression and Anxiety in People with Epilepsy. J Clin Neurol. 2014 Jul;10(3):175–188.

30 Pais, M.L., et al., Sex-specific cortical networks drive social behavior differences in an autism spectrum disorder model. Translational Psychiatry, 2025. 15(1): p. 251.

31 Reddy, D.S., W. Thompson, and G. Calderara, Molecular mechanisms of sex differences in epilepsy and seizure susceptibility in chemical, genetic and acquired epileptogenesis. Neuroscience Letters, 2021. 750: p. 135753.

32. Paolo Moretti, J. Adriaan Bouwknecht, Ryan Teague, Richard Paylor, Huda Y. Zoghbi, Abnormalities of social interactions and home-cage behavior in a mouse model of Rett syndrome, Human Molecular Genetics, Volume 14, Issue 2, 15 January 2005, Pages 205–220.

