## Supplementary figure and information for "Age dependent impairment of home cage behavior and reactivity in *Cntnap2* knock out mouse model"

### Content:

This supplementary information contains three supplementary figures.

**Supplementary Figure 1** – related to Figure 2, percentage nestlets torned

**Supplementary Figure 2** – related to Figure 5, comparison of freezing episodes

**Supplementary Figure 3** – related to Figures 6 and 7, morphological (qualitative) change in interneuron and microglia.

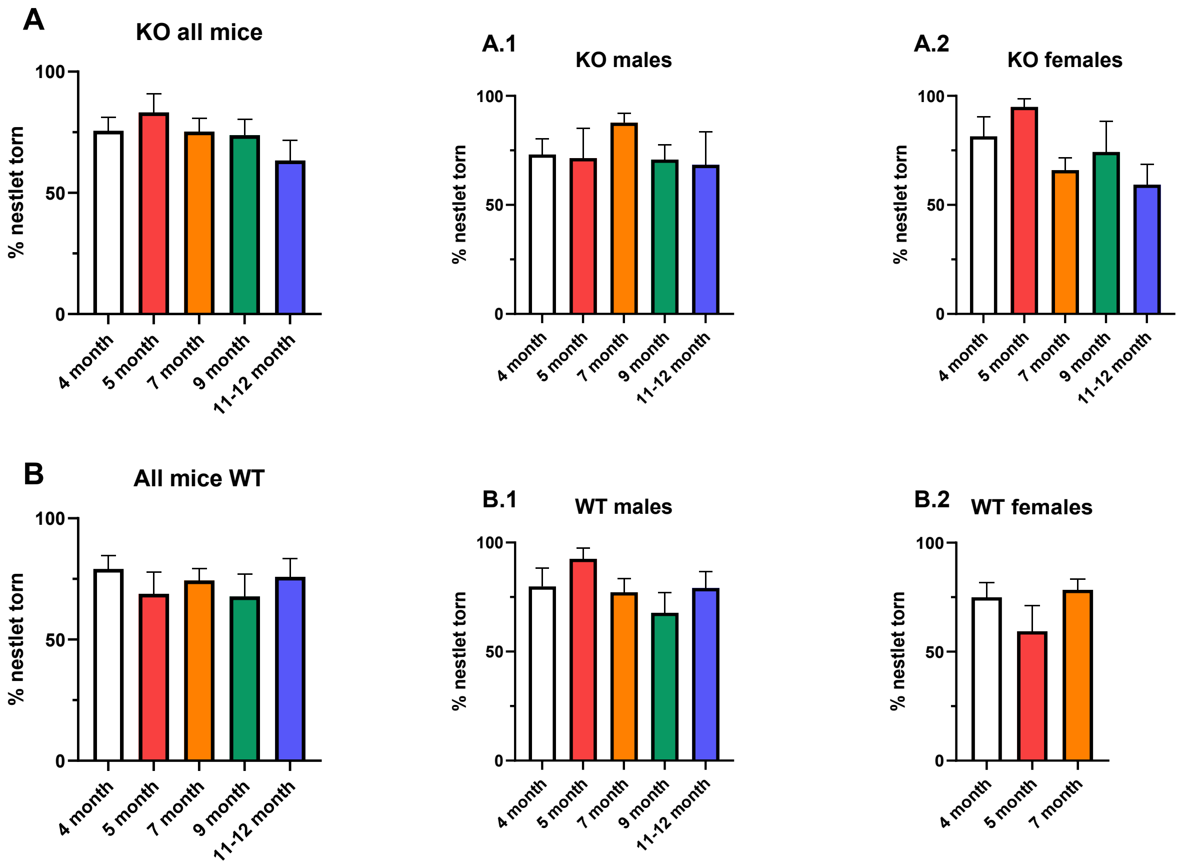


**Supplementary Figure S1.** Percent nestlet torn. (A) All KO mice did not have significant differences in % nestlet torn between any age groups, nor did KO males (A.1) or females (A.2) (Tukey's test, p>0.05, n=3-23). (B) All WT mice did not have significant differences in % nestlet torn between any age groups, nor did WT males (B.1) or females (B.2) (Tukey's test, p>0.05, n=6-15). There were no 9- or 11-12-month-old females.

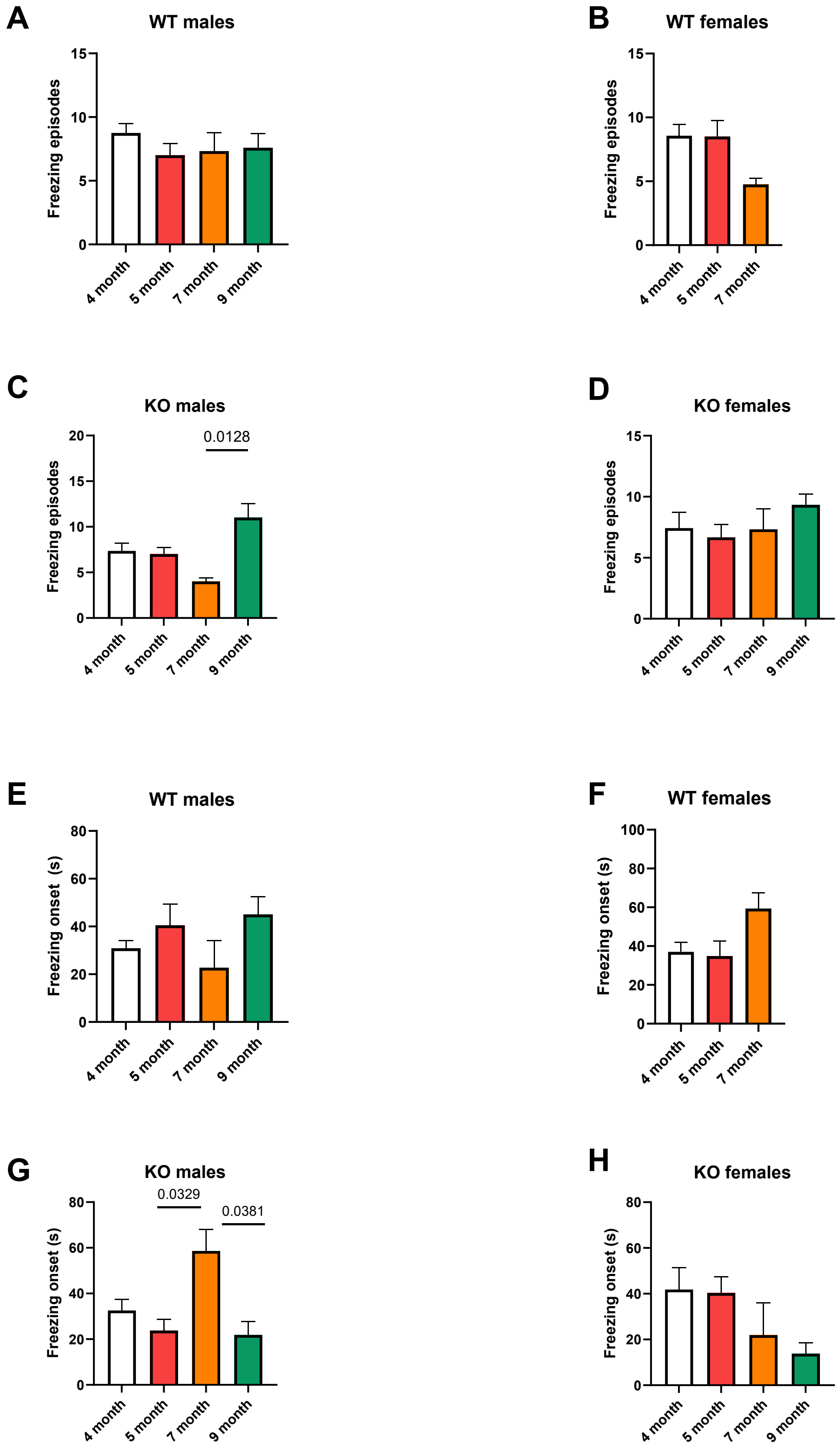
A
WT males
Freezing episodes
15
10
5
0
4 month
5 month
7 month
9 month
B
WT females
Freezing episodes
4 month
5 month
7 month
C
KO males
0.0128
20
D
KO females
E
WT males
Freezing onset (s)
80
60
40
20
F
WT females
100
G
KO males
0.0329
0.0381
H
KO females

**Supplementary Figure S2.** Sex-specific age-dependent freezing behavior in WT and Cntnap2 KO mice. (A-B) No age-dependent differences in freezing episodes were seen in male (A) and female (B) WT mice (Tukey's test, $p>0.05$, n=3-14). (C-D) 9-month-old Cntnap2 KO male mice (C) had a significantly increased number of freezing episodes compared to 7-month-old mice (Tukey's test, *$p=0.0128$, n=3, 4). No other age-dependent differences were seen in this cohort. In contrast, no significant age-dependent differences in number of freezing episodes were seen in KO female mice (D) (Tukey's test, $p>0.05$, n=3-7). (E-F) No age-dependent differences in freezing onset were seen in male (E) and female (F) WT mice (Tukey's test, $p>0.05$, n=3-14). (G-H) 7-month-old male Cntnap2 KO mice had a significantly later freezing onset than the 5 month ($p = 0.0329$) and 9-month groups ($p = 0.0381$) (G) (Tukey's test, n=4, 4, 3), while no significant age-dependent differences were found in female KO mice (H).

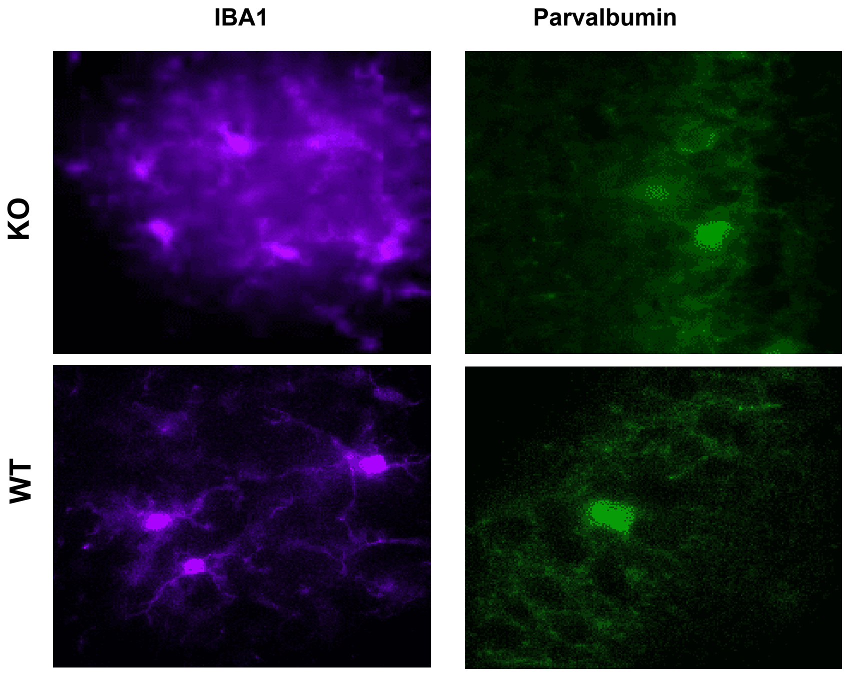


**Supplementary Figure S3:** (A) Representative image of parvalbumin (interneuron) and (B) IBa1 (microglia) immunostaining in *Cntnap2* mice and WT controls. An increased branching and size of microglia cells (cell body morphology) and an increased size of parvalbumin positive cells (morphology) was observed in hippocampus of *Cntnap2 KO* mice compared to the WT controls (qualitative analysis).
